# Association between mosaic loss of chromosome Y and pulmonary fibrosis susceptibility and severity

**DOI:** 10.1101/2024.05.25.595885

**Authors:** Dapeng Wang, Niran Hadad, Samuel Moss, Elena Lopez-Jimenez, Simon R. Johnson, Toby M Maher, Philip L Molyneaux, Yajie Zhao, John R. B. Perry, Paul J. Wolters, Jonathan A. Kropski, R Gisli Jenkins, Nicholas E. Banovich, Iain Stewart

## Abstract

Background

Pulmonary fibrosis (PF) is a rare lung disease with diverse pathogenesis and multiple interconnected underlying biological mechanisms. Mosaic loss of chromosome Y (mLOY) is one of the most common forms of acquired chromosome abnormality in men, which has been reported to be associated with increased risk of various chronic progressive diseases including fibrotic diseases. However, the exact role of mLOY in the development of PF remains elusive and to be elucidated.

**Methods:** We adopted three complementary approaches to explore the role of mLOY in the pathogenesis of PF. We used copy number on chromosome Y to estimate mLOY comparing patients in PROFILE and gnomAD cohorts and between cases and control patients from the GE100KGP cohort. Correlation of mLOY with demographic and clinical variables was tested using patients from PROFILE cohort. Lung single-cell transcriptomic data were analysed to assess the cell types implicated in mLOY. We performed Mendelian randomisation to examine the causal relationship between mLOY, IPF, and telomere length.

**Results:** The genetic analysis suggests that mLOY is found in PF from both case cohorts but when compared with an age matched population the effect is minimal (P = 0.0032). mLOY is related to age (P = 0.00021) and shorter telomere length (P = 0.0081) rather than PF severity or progression. Single-cell analysis indicates that mLOY appears to be found primarily in immune cells and appears to be related to presence and severity of fibrosis. Mendelian randomisation demonstrates that mLOY is not on the causal pathway for IPF, but partial evidence supports that telomere shortening is on the causal pathway for mLOY.

**Conclusion:** Our study confirms the existence of mLOY in PF patients and suggests that mLOY is not a major driver of IPF. The combined evidence suggests a triangulation model where telomere shortening leads to both IPF and mLOY.

## Introduction

Pulmonary fibrosis (PF) is a lung disease that is more prevalent in aging populations with particular genetic variants and its mortality rate is worse than most forms of the cancers (1). Environmental factors, smoking history and exposure to occupational hazards can increase the risk of the disease (2). Progressive loss of lung function caused by the lung scarring severely reduces the quality of life of the patients and the two frontline antifibrotic medications can only slow down the progression of the disease (3). The current understanding of the pathogenesis of pulmonary fibrosis highlights abnormal extracellular matrix deposition and remodelling during the repairing process in response to lung epithelial cell injury, followed by the activation of fibroblasts and interaction with other cell types through a number of signalling pathways (4–7).

Large-scale population based genetic analysis and genome-wide-association-studies (GWAS) provide an opportunity to identify susceptible genes and biomarkers for disease diagnosis and targeted therapeutics. Among the significant genes associated with PF, several genes play crucial regulatory roles in spindle assembly checkpoint (SAC) and proper chromosome segregation, such as *SPDL1* (8), *KIF15* (9), *MAD1L1* (9), and *KNL1* (10). Hypothetically, impairment and disruption of spindle assembly checkpoint mechanism could lead to ploidy errors and aneuploidy (11).

Mosaic loss of chromosome Y (mLOY), the most common form of aneuploidy relating to genome instability in non-malignant diseases, can occur in all age groups (12) but is most prevalent in older males. Smoking is a potential risk factor for mLOY in blood cells (13). GWAS variants significantly associated with mLOY are enriched among genes implicated in cell cycle regulation, hematologic malignancies and cancer susceptibility, suggesting the shared genetic architecture between mLOY and cancer susceptibility (14–16). mLOY increases the risk of prostate cancer (17), and mLOY is on the causal pathway of increased aspartate transaminase (AST) (18). Survival analysis reveals the association of mLOY in blood and the increased risk of all-cause mortality and non-haematological cancer mortality (19). mLOY is enriched in different cell types among different diseases, such as microglia in Alzheimer’s disease (20), bone marrow cells in myeloid neoplasm (21), and injured proximal tubule cells in chronic kidney disease (22). Experimentally created mLOY in murine hematopoietic stem and progenitor cells can lead to DNA damage, clonal hematopoiesis and acute myeloid leukemia (23).

It is reported that mLOY in blood is associated with cardiac dysfunction and fibrotic phenotype in various organs in a mouse model (24) and mLOY in blood cells profoundly contributes to the formation of cardiac fibrosis and the high mortality of patients after transcatheter aortic valve replacement via the activation of TGF-β signalling pathway (25). Although there is some evidence that mLOY might contribute to fibrogenesis, it is unclear whether mLOY is directly causally associated with PF or its severity. We therefore hypothesised that the genetic variants associated with IPF would promote fibrogenesis via mLOY. To test this hypothesis, we assessed the copy number on chromosome Y in two case-control cohorts and the correlation between mLOY and demographic/clinical variables in PROFILE cohort. Secondly, we examined cell types that could be driving the difference between disease and control using single-cell transcriptomic data. Thirdly, we conducted Mendelian randomisation between mLOY, IPF and telomere length.

## Methods

### Cohorts

The whole genome sequencing data from two independent PF case cohorts were used in the study: PROFILE cohort and GE100KGP cohort. PROFILE is a UK based cohort of treatment naive PF patients recruited from two locations including London and Nottingham in the United Kingdom (8). This cohort comes with a rich set of demographic and clinical baseline and longitudinal meta data (26). GE100KGP case cohort is defined based on the health record data and/or clinically diagnosed familial pulmonary fibrosis diseases from the 100K Genome Project run and managed by Genomics England, which has been described previously. For each cohort, only male patients were used for the analysis of mLOY. Sample metadata for samples in the HGDP and 1KG subsets were downloaded from gnomAD (v3.1.2) as controls for PROFILE (27). To mitigate the confounding effect of age, propensity score matching was performed to generate the non-PF controls in the rare disease programme from 100K Genome Project data to match the case cohort based on age information (28).

### Estimate of copy number

The male-specific region Y (MSY) is unique to chromosome Y and not shared by chromosome X. Therefore, MSY was chosen for analysis. The copy number of chromosome Y was defined as the mean read depth on MSY divided by mean read depth on 22 autosomes, unless otherwise stated. The read depth was calculated using SAMtools from the BAM files of whole genome sequencing data mapped onto human reference genome (GRCh38) (29). Due to data availability, a different approach was used to measure copy number on chromosome Y for samples from gnomAD cohort: copy number on chromosome Y was defined as the sample’s mean depth across chromosome Y divided by sample’s mean depth across chromosome 20. Patients in PROFILE cohort were divided into mLOY group and non-mLOY group where mLOY group is defined as those patients with less than mean minus standard deviation of copy number on chromosome Y.

### Correlation analysis with clinical and demographic variables in PROFILE cohort

Five continuous variables including age, ppFVC (lung volume), ppTLCO (gas transfer), CPI (composite physiological index), telomere length and two discrete variables including gender age physiology stage and disease progression in 12 months were selected to study their correlation with mLOY where disease progression was defined as a 10% relative decline in FVC or death within 12 months from the baseline visit and disease progression was clinically adjudicated when lung function follow-up was missing but participants survived beyond 12 months. Fisher’s exact test was used for testing the relationship between mLOY and risk alleles from four spindle assembly checkpoint (SAC) genes. Wilcoxon test and Kruskal-Wallis test were used for testing between two groups and three groups, respectively. Spearman’s rank correlation coefficient was used for the correlation analysis between two variables. The significance of the tests was determined at P < 0.05.

### Single-cell analysis

Single-cell analysis was carried out using 100 samples collected from 69 male patients with either PF or controls (Control = 30, IPF = 22, ILD = 5, CTD-ILD = 1, CWP = 3, sarcoidosis = 2, NSIP = 2, IPAF = 2 and cHP = 2) from a previously published single-cell RNA-sequencing dataset (GSE227136; (30)). Data were collapsed in cases where multiple samples were collected from the same patient, intended to represent less and more fibrotic regions, unless specified otherwise. CellRanger output BAM files were used to extract reads uniquely mapped to chromosome Y and chromosome 12. Per cell mLOY was determined as the ratio between reads mapped to Y and reads mapped to chr12, limiting the analysis to cells that passed filtering using the filtering criteria described in (30). Cells with chrY/chr12 ratio equal to 0 were considered cells with mLOY. The proportion of mLOY for a sample was determined as the proportions of cells without reads mapped to Y (mLOY = 0).

To assess the probability of mLOY between disease and control, and with disease progression (less fibrotic compared with more fibrotic regions) cells were assigned to a binary mLOY positive or mLOY negative category and logistic regression model was fitted for each of the following covariates: disease type, lineage and cell identity. Significance was set to FDR adjusted p-value < 0.05.

Differential expression was assessed between mLOY cells and the top 25th percentile of non-mLOY cells in disease samples using negative binomial mixed models in both cell-agnostic and cell-type aware manner. A model was fitted for each cell type assessing changes to gene expression associated with mLOY using the *glmmTMB* package (31) with a negative binomial distribution and a log link function. Sample identity was used as random effect to account for dispersion associated with single-cell data. Y-chromosome genes were excluded from the analysis, and genes with FDR-adjusted P<0.05 were considered as significant. Standardized regression coefficients were obtained using the *effectsize* package (32) and used for gene set enrichment analysis (GSEA) using the *clusterProfiler* package (33).

### Mendelian randomisation

This two sample MR approach used public GWAS summary statistics to identify genetic instruments (single nucleotide polymorphisms; SNPs) for IPF susceptibility (4125 cases, 20 464 controls) (10), ovarian cancer (25 509 cases, 40 941 controls) (34), prostate cancer (79 148 cases, 61 106 controls) (35), and telomere length (leukocyte) (sample size: 472 174) (36). Summary statistics for mLOY were retrieved upon request from authors JRBP and YZ (sample size: 204 770).

Genetic instruments were filtered for genome-wide significance (p < 5×10^-8^) and clumped to ensure independence using the 1000 Genomes Project reference panel (r2 < 0.001 within a 10,000 kb window). All SNPs were required to be strong instruments for each exposure (F > 10) to avoid weak instrument bias.

A primary bidirectional MR analysis was performed to assess for causal effects between mLOY and IPF. A random-effects inverse-weighted (IVW-RE) MR method (37) was used to provide main causal estimates (*MendelianRandomization* (38)). MR-Egger (39) and weighted median (40) methods were used to provide pleiotropy-robust MR estimates. MR-PRESSO (41) and MR-Lasso (42) methods were used to provide pleiotropy-robust MR estimates by identifying and removing genetic instruments with likely horizontal pleiotropic effects. A SNP leave-one-out analysis was performed to identify SNPs that are independently driving or distorting causal effect estimates. As mLOY is a sex-specific trait, a negative control analysis assessing effects of the mLOY instrument on ovarian cancer risk was performed to evaluate gene-environment equivalence. A positive control analysis also assessed the effects of the mLOY instrument on prostate cancer risk. As shortened telomere length is a risk factor for IPF, a secondary analysis was performed to assess if telomere length may also have causal effects on mLOY.

## Results

### mLOY is observed in PF

In total, 387 male patients in PROFILE case cohort and 232 male patients in GE100KGP case cohort as well as 2216 male patients in gnomAD control cohort and 232 male patients in GE100KGP control cohort were included. The copy number on chromosome Y was significantly lower in PROFILE case cohort (median: 0.309) than gnomAD control cohort (median: 0.375; P < 2.22e-16) (Figure 1A). To account for age related effects a propensity matched non-PF control cohort was generated from the GE100KGP dataset, and the copy number on chromosome Y in GE100KGP case cohort (median: 0.288) was marginally but significantly lower than that in GE100KGP control cohort (median: 0.291; P = 0.0032) (Figure 1B). Overall, lower copy number on chromosome Y was observed in PF patients compared with non-PF patients in both cohort analyses, indicating an association between mLOY and PF. However, age was highly negatively correlated with copy number on chromosome Y for both PROFILE cohort (Spearman: R = -0.24, P = 1.2e-6) and GE100KGP cohort (Spearman: R = -0.38, P = 2.3e-9).

**Figure 1.**
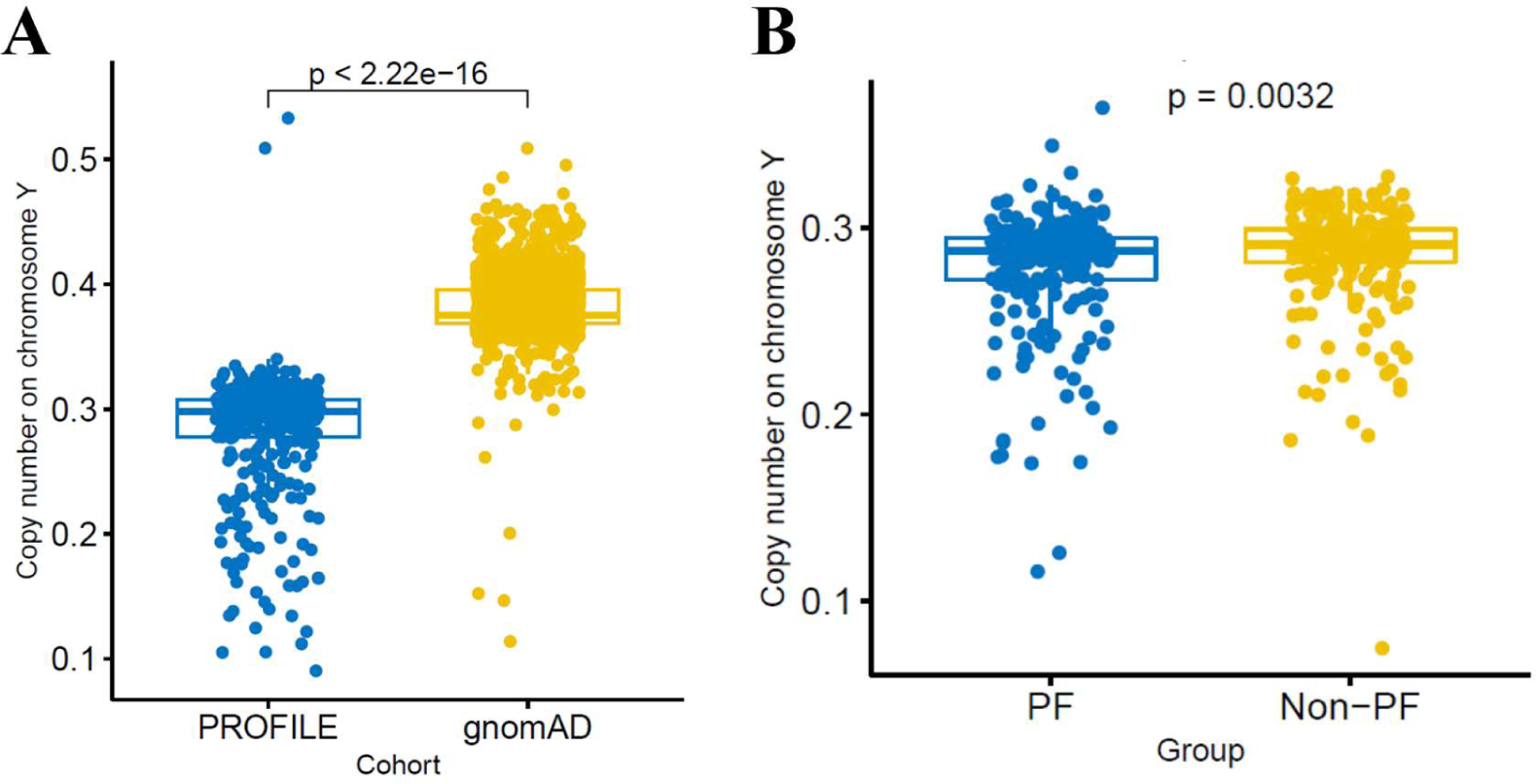
mLOY in case and control cohort. (A) Copy number on chromosome Y for male PF case from PROFILE and non-PF control patients from gnomAD cohort. Copy number on chromosome Y is defined as mean read depth on chromosome Y divided by that on chromosome 20. (B) Copy number on chromosome Y for male PF case and non-PF control patients from GE100KGP cohort. Copy number on chromosome Y is defined as mean read depth on MSY divided by mean read depth on 22 autosomes. Wilcoxon test was used for testing the difference between two groups.

### mLOY is related to age and telomere length but not PF severity or disease progression

To investigate whether mLOY is directly involved in PF pathogenesis and the relationship between mLOY and key demographic features, we defined a threshold for substantial mLOY as a copy number of <0.2565 and non-mLOY as a copy number of >=0.2565. Patients with substantial mLOY were significantly older (75 vs 71; P = 0.00021) and had significantly shorter telomeres (0.728 vs 0.775; P = 0.0081) than patients without mLOY. However, no significant difference in ppFVC (P = 0.63), and ppTLCO (P = 0.75) were found (Figure 2). Similarly, correlation analysis found no association between the lung function measurements and copy number on chromosome Y: ppFVC (R = -0.023; P = 0.65), and ppTLCO (R = 0.036; P = 0.5) (Figure S2). Furthermore, no significant difference was found in the copy number on chromosome Y between patients that did progress in 12 months (N = 179) and patients that did not progress in 12 months (N = 208; P = 0.55) (Figure 3). Stratification of mLOY by the risk alleles from SAC genes associated with IPF did not enrich patients with mLOY suggesting that the causal mechanisms for SAC genes was not directly related to mLOY (Figure S3).

**Figure 2.**
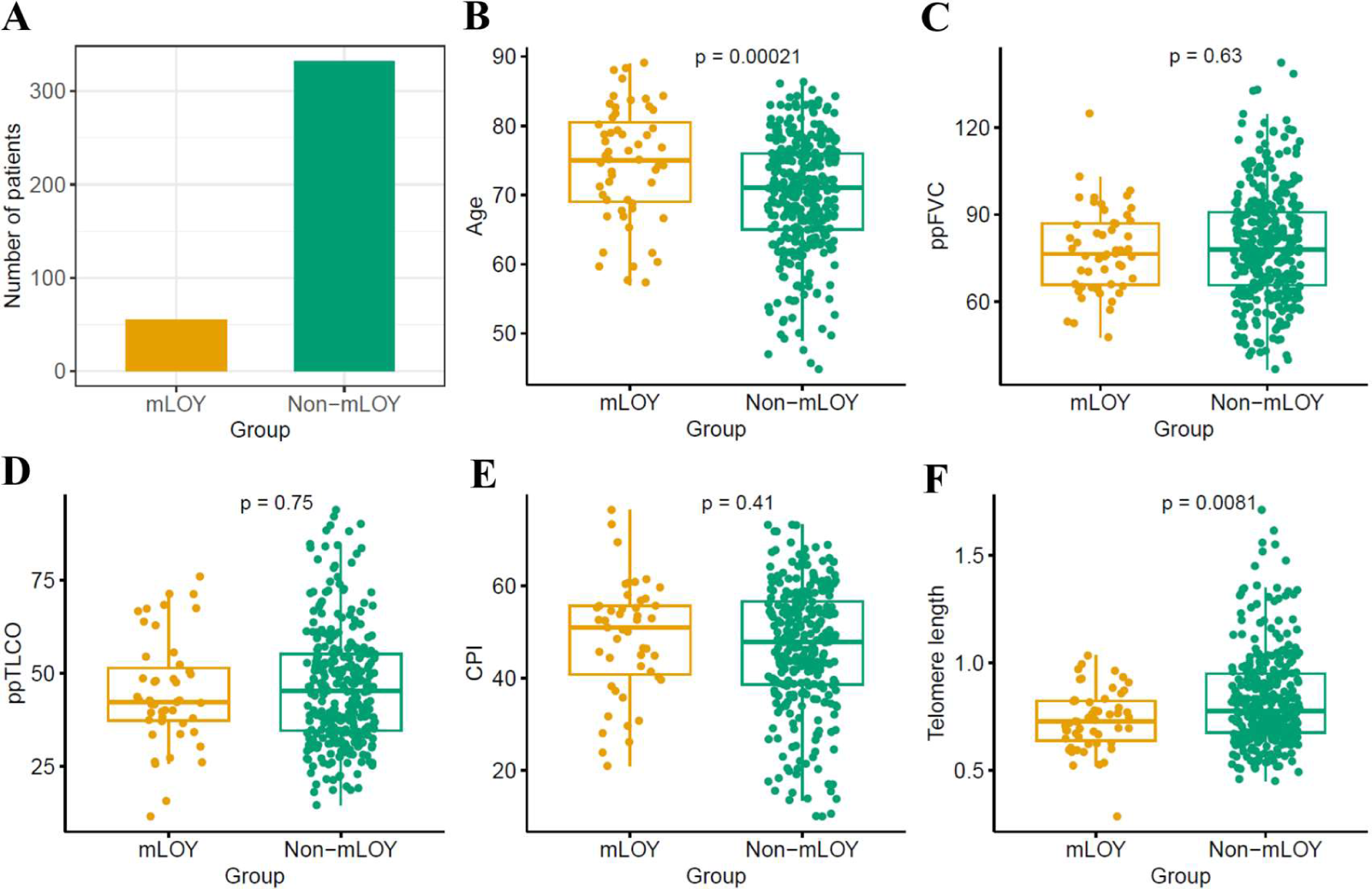
Compare the age, lung function and telomere length between mLOY and non-mLOY group male PF patients from PROFILE cohort. (A) Number of PF patients of mLOY and non-mLOY group (B) Age (C) ppFVC (D) ppTLCO (E) CPI (F) Telomere length. Wilcoxon test was used for testing the difference between two groups.

**Figure 3.**
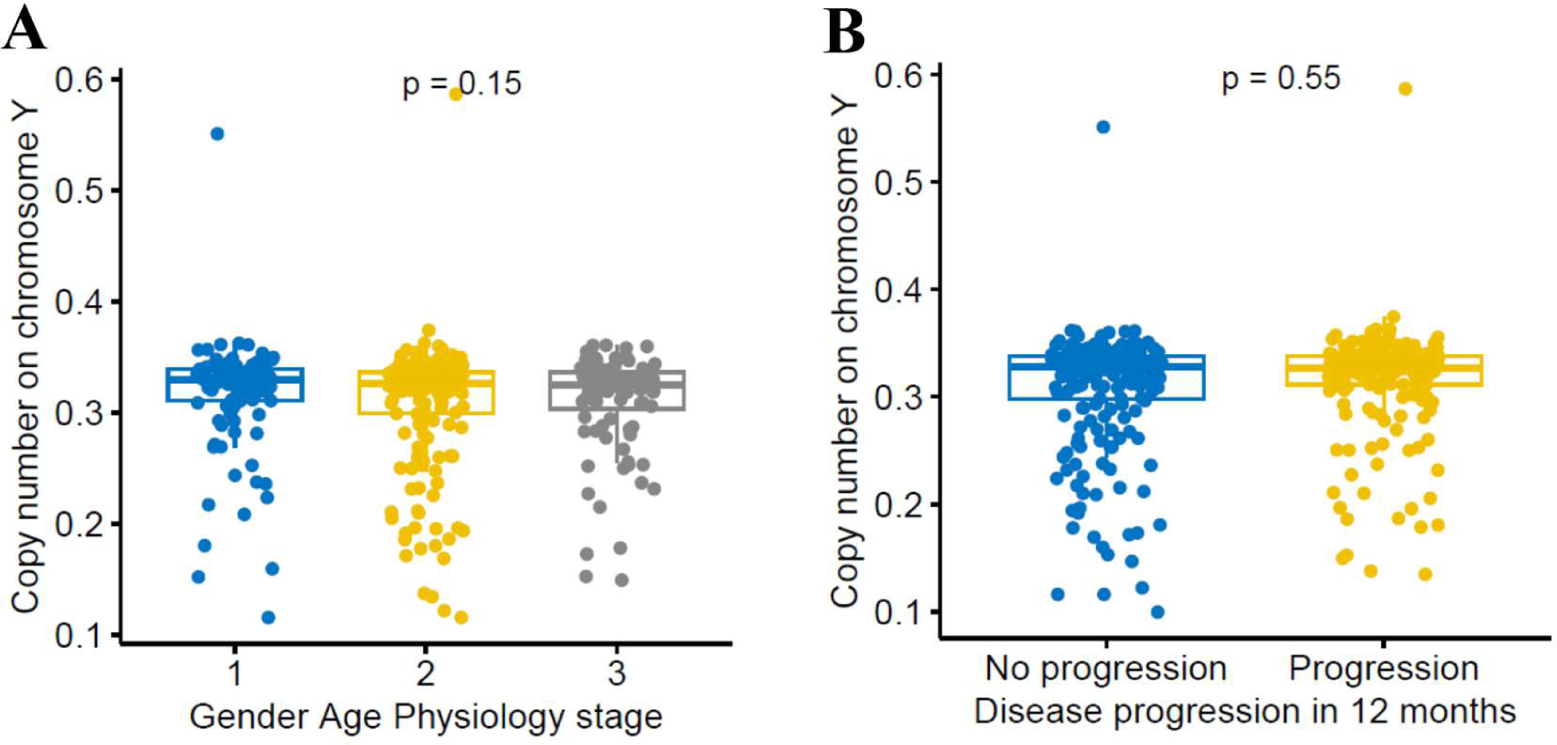
Correlation of mLOY for male PF patients from PROFILE cohort with disease severity. (A) Gender Age Physiology stage. Kruskal-Wallis test was used for testing the difference among the three groups. (B) Progression in 12 months. Wilcoxon test was used for testing the difference between two groups. Copy number on chromosome Y is defined as mean read depth on MSY divided by mean read depth on 22 autosomes.

### Single-cell analysis

To understand whether mLOY in structural cells is associated with pulmonary fibrosis, mLOY was assessed using lung single-cell transcriptomic data from 39 males with PF and 30 control males. Cells from PF samples have a higher probability to lose chromosome Y compared with controls, as evidenced by the absence of mapped read to chromosome Y (Figure 4, Figure S4A). Among the different types of ILDs, sarcoidosis, cHP, NSIP, CWP, IPF, CTD-ILD and other ILDs show significantly higher probability of mLOY compared with controls (Figure 4B, Figure S4B). To elucidate whether mLOY exhibits cell-type specificity, the probability of mLOY occurring in a cell-type specific manner was assessed. Cells of immune lineage had significantly higher probability for mLOY in fibrotic tissue compared with controls (Figure 5A), including dendritic cells, CD4+ and CD8+ cells, monocytes, and alveolar macrophages. Among PF subjects who had paired samples from more and less fibrotic regions of the lung, alveolar macrophages from more fibrotic regions had a higher probability of mLOY compared to samples from less fibrotic regions (Figure 5A). mLOY in other immune cells showed a similar trend (Figure 5B). In contrast with immune cells, epithelial cells showed lower probability of mLOY, except for AT2 cells which had significantly higher probability for mLOY in fibrotic tissue (Figure 5A).

**Figure 4.**
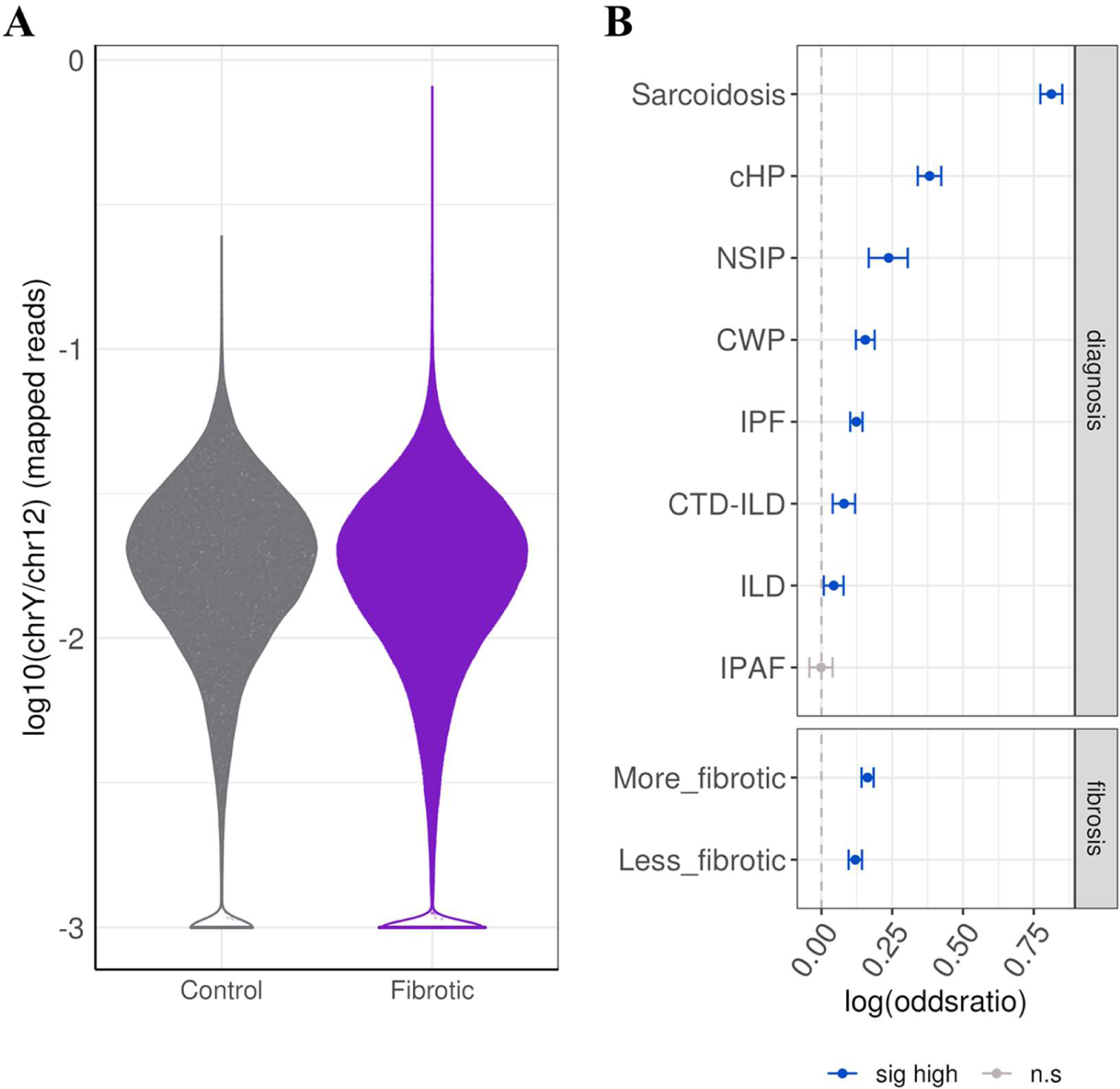
mLOY increases with disease severity. (A) Distribution of chrY/chr12 ratio across individual cells collapsed by control aged samples and fibrotic samples. Plot is scaled by width. (B) Association between mLOY probability and disease according to ILD type and fibrosis status. Significant results (FDR adjusted p-value < 0.05, logistic regression) are coloured in blue. Values represent odds ratio and 95% CI.

**Figure 5.**
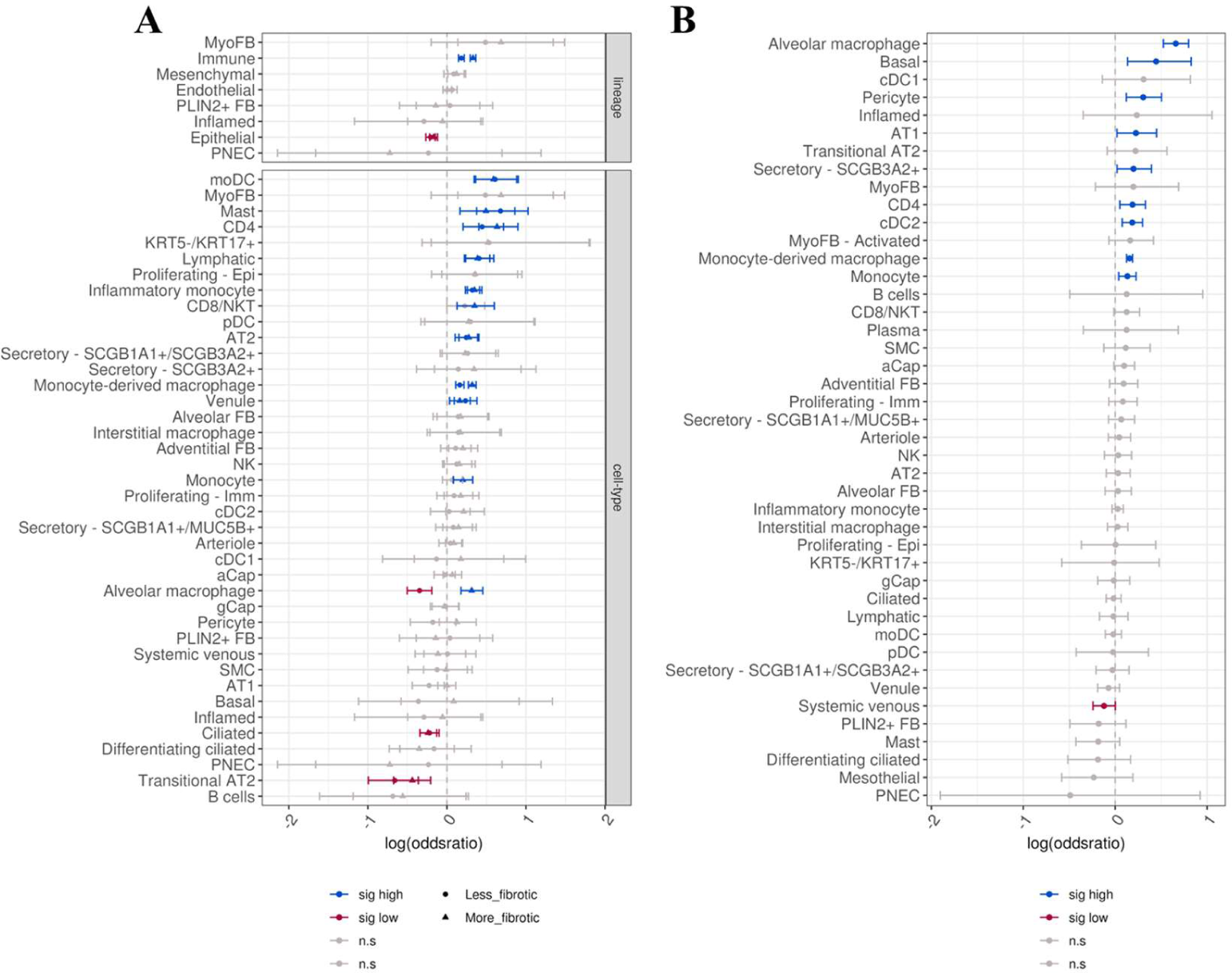
mLOY increases with fibrosis primarily in immune cells. (A) Association between mLOY probability and disease according to disease or cell-type. Shape corresponds to less fibrotic (circle) or more fibrotic (triangle) samples. (B) Association between mLOY probability and disease progression comparing more fibrotic samples to less fibrotic samples. Significant results (FDR adjusted p-value < 0.05, logistic regression) are coloured by either blue or red, highlighting higher or lower probability of mLOY with fibrosis, respectively. Values represent odds ratio and 95% CI.

To assess whether mLOY is associated with changes in expression signature, differential expression analysis was carried out using cell-type agnostic and cell-type specific approaches between mLOY cells and the top 25th percentile of cells based on their chrY/chr12 ratio. Differential expression by cell-type showed that expression changes associated with mLOY were predominately observed in immune cells, including monocyte-derived macrophages, inflammatory monocytes, CD4+ cells, and proliferating immune cells (Figure S5A). However, high number of differentially expressed genes were also observed in epithelial cells (ciliated cells, MUC5B+ secretory cells), endothelial cells (systemic venous, venule) and mesenchymal cells (alveolar fibroblasts). Generally, expression changes were cell-type specific and show little overlap across cell types (Figure S5B). Gene set enrichment analysis shows that in immune cells mLOY was associated with activation of genes of metabolic pathways (Figure S5C). To assess cell-common gene expression changes that are associated with mLOY, differential expression was performed using a cell-type agnostic approach. A total of 1,399 differentially expressed genes were identified (Figure S5A, FDR adjusted P < 0.05 & |std. regression coefficient| > 0.05). Enrichment analysis reveals activation of genes related to the lysosomal and phagosome, activation of immune-related signalling pathways and apoptosis (Figure S5C). The single-cell transcriptomic data support the role of specific types of immune cells in mLOY in PF patients.

### mLOY is not on the causal pathway for PF but short telomeres are on the causal pathway for mLOY

To understand the causal relationship between mLOY, PF and telomere length, Mendelian randomisation was performed. For Mendelian randomisation between mLOY and IPF, a negative effect of mLOY on IPF was identified (IVW-RE, OR = 0.526; 95%CI = 0.328-0.845; P = 0.008) through the IVW-RE method, however, no effects were detected after correcting for pleiotropy (Figure 6A). No causal effect of IPF on mLOY was identified in all five methods (IVW-RE, OR = 0.997; 95%CI = 0.989-1.004; P = 0.377) (Figure 6B). As a negative control, no causal effect of mLOY on ovarian cancer was found for all five methods (IVW-RE, OR = 1.047; 95%CI = 0.865-1.269; P = 0.634) (Figure S6A) and as a positive control, causal effect of mLOY on prostate cancer was identified from all five methods (IVW-RE, OR = 1.461; 95%CI = 1.247-1.711; P = 2.56E-06) (Figure S6B), which is consistent with the effects identified in a previous study (14). Three out of five methods showed the protective causal effect of telomere length on mLOY although the result from IVW-RE method was not significant (IVW-RE, OR = 0.963; 95%CI = 0.915-1.014; P = 0.153) (Figure 6C). Furthermore, leave-one-out sensitivity analyses identified significant effects of shorter telomere length on mLOY when either rs16978028 or rs7705526 is removed (Figure S7).

**Figure 6.**
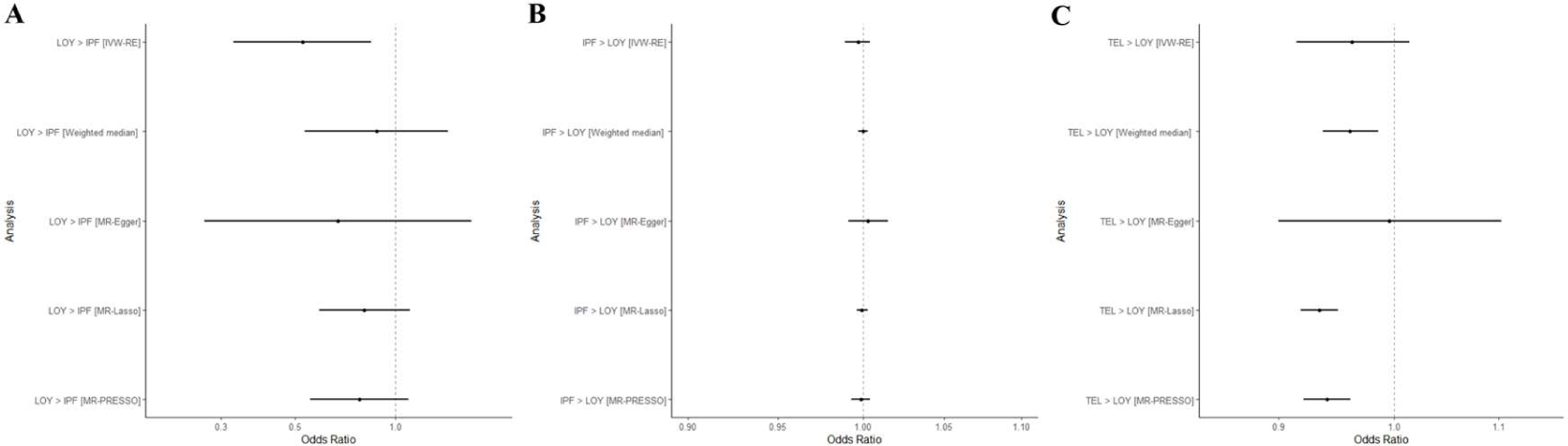
Forest plot from Mendelian randomization analysis for the causal association. (A) mLOY and IPF (B) IPF and mLOY (C) Telomere length and mLOY.

## Discussion

These data demonstrate that mLOY is greater in male patients with PF compared with non-PF patients, however, peripheral blood mLOY is likely to be driven in large part by age and telomere length, and there was no association with disease severity or progression. Single-cell transcriptomic analysis of lung tissue from patients with a variety of fibrotic diseases identified that mLOY in immune cells is associated with fibrotic diseases and there appeared to be an association with increased fibrosis within samples. However, bidirectional Mendelian randomisation did not find any causal effects, but there was suggestive evidence that shorter telomere length was on the causal pathway for mLOY.

mLOY and PF share several risk factors such as age, smoking and telomere shortening. It is well known that the positive correlation exists between the prevalence of mLOY and age in the general male population (14). The subtle difference in mLOY between PF case cohort and age-matched non-PF cohort suggests that effect of age is driving mLOY in IPF. Previous Mendelian randomisation analysis has demonstrated that telomere shortening is on the causal pathway for IPF but not COPD (43). The indication that telomere length is on the causal pathway of both IPF and mLOY supports a model where telomere length is on the causal pathway for both IPF and mLOY which may explain some co-morbidities more directly attributable to mLOY (24).

It has been widely recognised that a variety of innate and adaptive immune cells interacting with other types of cells could be involved throughout the development of IPF, especially after lung injury and infection (44). A feed forward mechanism suggests that activation of immune cells and fibroblasts releases pro-inflammatory cytokines and chemokines to strengthen the existing activation (45). The direct causal relationship between immunophenotypes and IPF risk have been confirmed by a Mendelian randomization analysis (46). Dysfunctional immune system with other pro-fibrotic pathways may contribute to the long-term unresolved inflammation leading to the formation of fibrosis while different types of cells might play opposite roles, suggested by studies on IPF and mouse models (47). Our study identifies three types of immune cells in which mLOY is related to fibrosis disease and fibrosis severity: CD4 cells, monocyte-derived macrophages and alveolar macrophages. The number of CD4+ cells is reported to increase in IPF lungs, suggesting its functional relevance to IPF (48). CD4+ T lymphocytes have been found to be associated with increased level of mLOY in prostate cancer patients whereas mLOY is enriched in NK cells for Alzheimer’s disease patients, suggesting that the association of CD4 cells with mLOY exhibits disease specific features (49). Macrophages represent the most common immune cells in the human lung and alveolar macrophages are one of the two major types of pulmonary macrophages (50). Significantly reduced efferocytosis in alveolar macrophages and impaired mitochondrial homeostasis in alveolar macrophages observed in IPF patients link alveolar macrophages to the pathogenesis of IPF. Alveolar macrophage can also impact normal and IPF fibroblasts to differing extents through cell-cell communication (51). Pro-inflammatory cytokine expressed classically activated (M1) and anti-inflammatory cytokine expressed alternatively activated (M2) phenotypes exert polarised effects on the responses to the injury (50). Persistent lung injury recruits the monocyte-derived macrophage cells and transits macrophages into M2 phenotype under the regulation of TGF-β signalling pathway (52, 53). The findings that these important immune cells experimentally proved to be involved in the IPF disease molecular mechanism are profoundly impacted by mLOY highlight the potential role of mLOY in the pathogenesis of IPF. This will provide valuable information for the appropriate design and use of treatments for IPF patients, such as immunosuppressive medications.

This study has a number of strengths including the use of propensity matching, analysis of single cells and the undertaking of Mendelian randomisation. For each type of analysis, we either performed the analysis on multiple case-control cohorts or conducted sensitivity analyses to generate the most robust results. However, there were several limitations. We used the copy number on chromosome Y from whole genome sequencing as a proxy for mLOY for each patient, and mLOY status could be confirmed using the experimental approaches such as fluorescence in situ hybridization (FISH) and droplet-digital PCR (ddPCR) in the future. The study is limited by the availability of the specific datasets. For the genetic analysis, GE100KGP cohort is a disease enriched cohort with which the analysis might overestimate the strength of mLOY in the age-matched control. For the future work, a case-control cohort study to include the PF cases and age-matched healthy controls is needed to validate the findings.

In conclusion, whole genome sequencing data analysis, single-cell transcriptomic analysis and Mendelian randomisation analysis show that mLOY is related to PF but mLOY is not on the causal pathway of PF. We propose that telomere shortening is causing both mLOY and PF through different molecular mechanisms. Further experimental validation should be performed to test this hypothesis.

## Acknowledgements

This work was funded by an MRC Programme Grant awarded to RGJ. This research was made possible through access to data in the National Genomic Research Library, which is managed by Genomics England Limited (a wholly owned company of the Department of Health and Social Care). The National Genomic Research Library holds data provided by patients and collected by the NHS as part of their care and data collected as part of their participation in research. The National Genomic Research Library is funded by the National Institute for Health Research and NHS England. The Wellcome Trust, Cancer Research UK and the Medical Research Council have also funded research infrastructure. Additional support was provided by the US National Institutes of Health (R01HL145372, NEB/JAK), the Department of Defense (NEB/JAK), and the Department of Veterans Affairs (JAK).

## Contributions

DW, PJW, JAK, RGJ, NEB and IS conceived and designed this study. DW, NH and SM conducted the data analysis. EL, SRJ, TMM, PLM, YZ, and JRBP contributed to the patient recruitment and data collection. DW wrote the paper with input from all authors.

