## Supplementary Material for "Association between mosaic loss of chromosome Y and pulmonary fibrosis susceptibility and severity"

### Supplementary information

#### Figure S1. Age distribution for male PF patients.

(A) PROFILE (B) GE100KGP cohort.

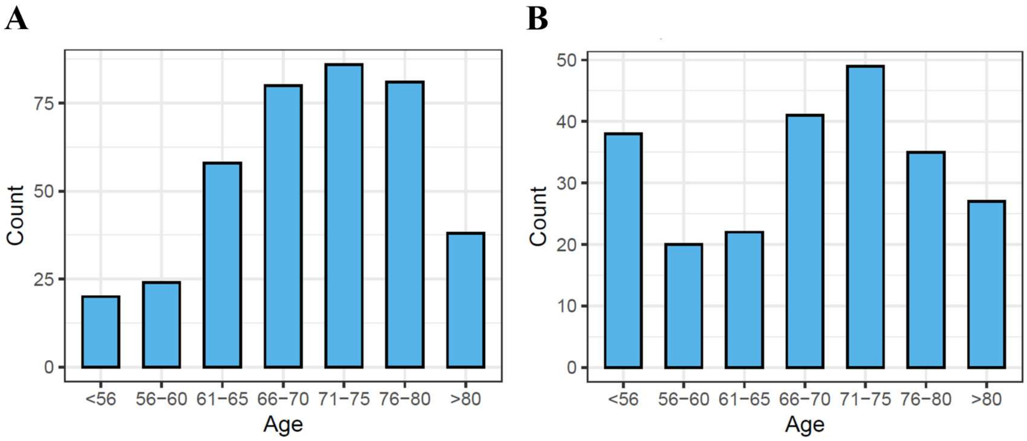

**Figure S2. Correlation between lung function and mLOY for male PF patients from PROFILE cohort.**

(A) ppFVC (B) ppTLCO (C) CPI (D) Telomere length. Copy number on chromosome Y is defined as mean read depth on MSY divided by mean read depth on 22 autosomes.

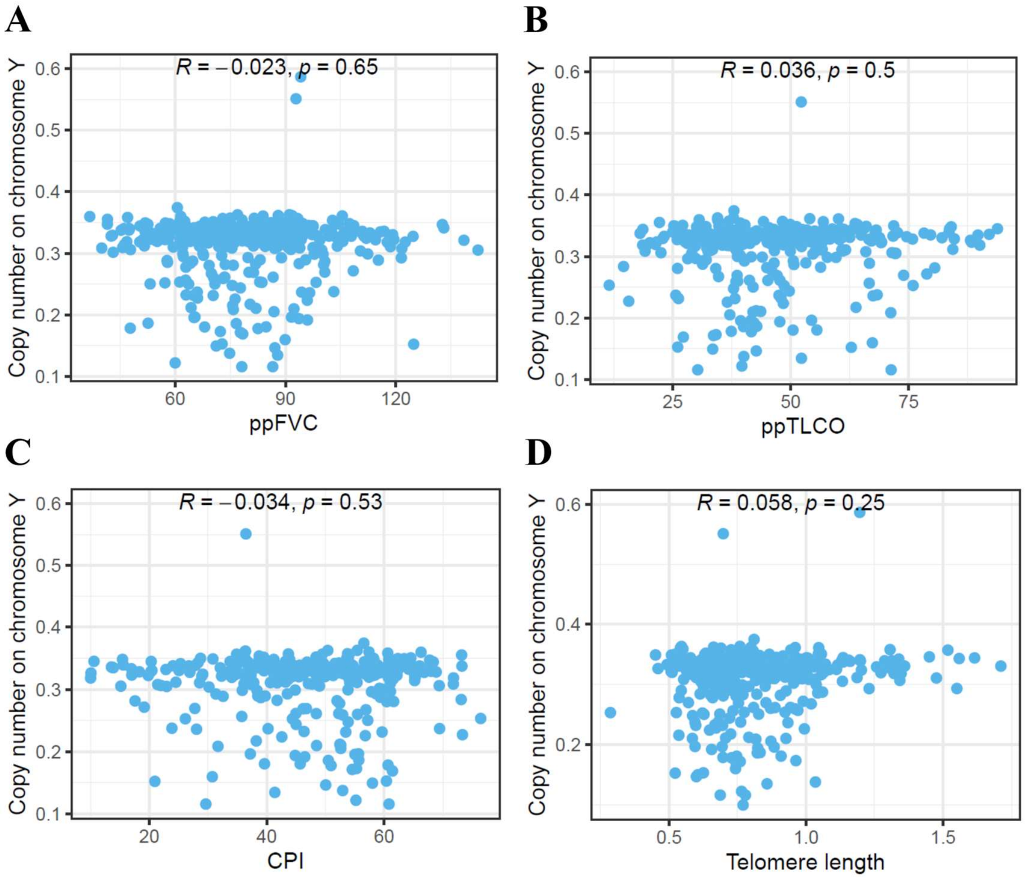

**Figure S3. Association of mLOY and risk alleles from four SAC genes for PF male patients from PROFILE cohort.**

(A) Number of patients without risk alleles and with risk alleles. (B) Fisher's exact test for relationship between mLOY and risk alleles.

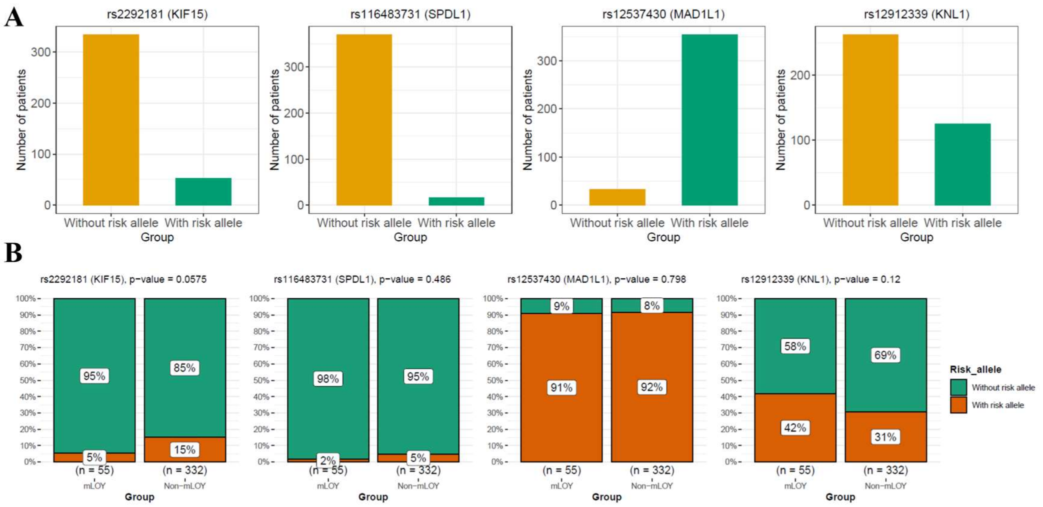

**Figure S4. mLOY across samples and disease types.**

Percent of cells showing mLOY across samples coloured by (A) fibrosis status and (B) disease status.

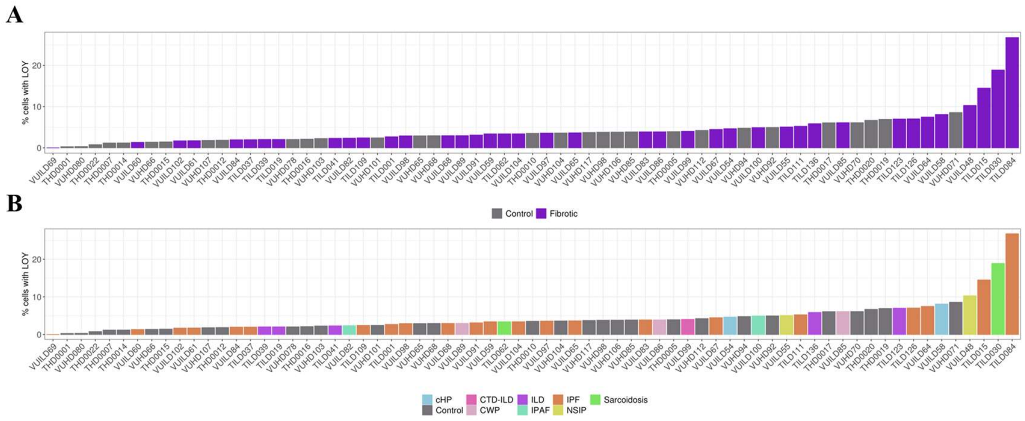

**Figure S5. Differential expression between cells lacking chromosome Y and top 25% Y/chr12 ratio.**

(A) Number of differentially expressed genes between cells with mLOY and the top 25% of cells for chrY/chr12 ratio (B) Upset plot showing the intersection between differentially expressed genes across cell types (C) GSEA results for each cell types.

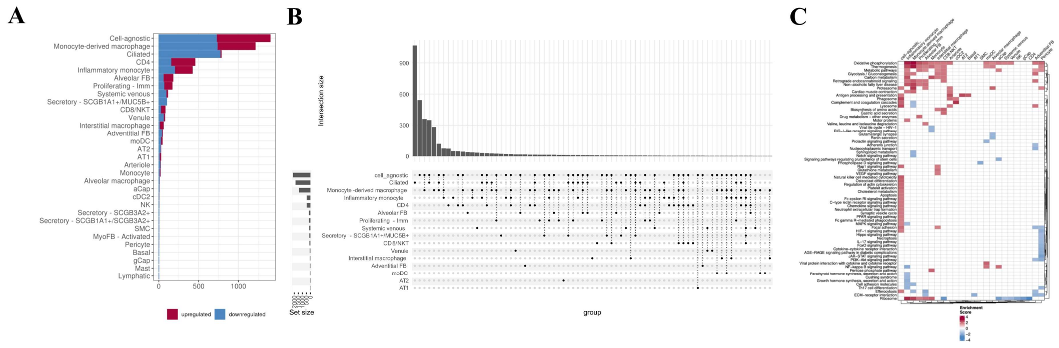

**Figure S6. Forest plot from Mendelian randomization analysis for the causal association.**

(A) mLOY and ovarian cancer (negative control) (B) mLOY and prostate cancer (positive control).

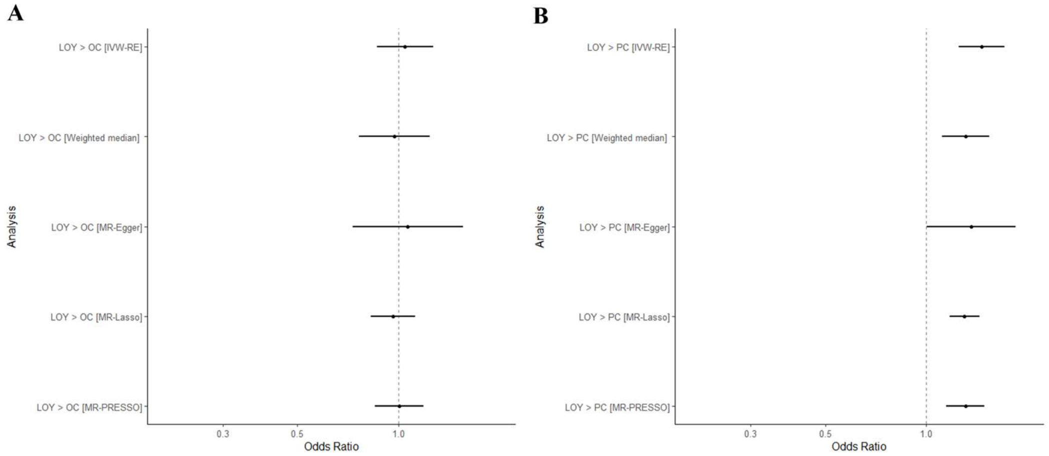

**Figure S7. Results of Mendelian randomization analysis for the causal association between telomere length and mLOY.**

(A) Scatter plot (B) Leave one out plot (C) Single-SNP estimates forest plot.

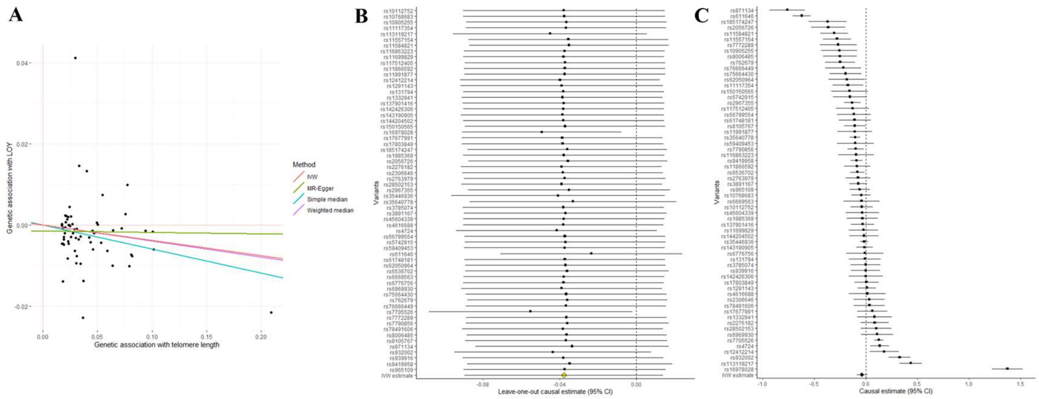
